# Expression and functional evaluation of recombinant human ABO blood group antigen cleaving glycoside hydrolases α-N-acetylgalactosaminidase, α-galactosidase, and endo-β-galactosidase produced in *Escherichia coli*

**DOI:** 10.1101/2022.05.04.490255

**Authors:** Wu Han Toh, Yvonne Kuo, Sean Kai Hsu, Bernie Chen, Alan Justin Lee, Easton Liaw, Jane Lee, Alexander Cheng, Laura Hwa, Kaitlyn Hu, Sienna Chien, Christine Wong, Kristin Chang, Minna Hang, Sabrina Hong, Ethan Su, Jude Clapper, Jonathan Hsu

**Author notes:** These authors contributed equally to this work (Wu Han Toh, Yvonne Kuo).

## Abstract

Blood transfusions are an integral component of healthcare; however, availability of viable blood is limited by patient-donor blood type specificity, which contributes to seasonal shortages as well as shortages worldwide, especially in developing countries, and during pandemics or natural disasters. Attempts to increase blood supply with commercial incentives have raised ethical concerns, and current proposed artificial blood substitutes are unable to fully replicate the function of native red blood cells (RBCs). In this study, we explore the potential strategy of alleviating blood shortages through enzymatic conversion of A, B, and AB blood types to blood type O. In theory, this process eliminates ABO patient-donor incompatibility, which increases the supply of universal donor blood. Three glycoside hydrolases, α-N-acetylgalactosaminidase, α-galactosidase, and endo-β-galactosidase, were selected to act as molecular scissors to cleave terminal residues on A and B RBC surface antigens and catalyze the conversion process. These enzymes were recombinantly expressed in BL21(DE3) *Escherichia coli* and purified through nickel ion affinity chromatography. A combination of colorimetric substrate assays, thin-layer chromatography, and mass spectroscopy were utilized to evaluate enzyme functionality. Enzyme efficiency was modeled using Michaelis–Menten kinetics. Partial enzymatic A-to-O blood type conversion on porcine red blood cells was observed with slide agglutination tests. Results confirm recombinant enzyme-mediated blood type conversion as a potential strategy for alleviating blood shortages.

## Introduction

Blood transfusions are an integral component of healthcare used to replace blood that is lost or provide blood when one’s body lacks normal production. They are needed in both routine and emergency situations, such as after an injury, during surgery, and for people with blood related disorders such as anemia, sickle cell disease, hemophilia, and cancer (NIH, 2022). Transfusions save millions of lives each year, drastically improving patients’ life expectancy and quality of life (WHO, 2010).

The availability of blood for transfusions is regulated by the supply and demand of safe, transfusable blood. Because blood products can only be derived from healthy donors, supply is limited by public willingness to donate. In the past, the WHO has estimated that a minimum donation rate of 1% of the total population is required to satisfy the most basic requirements for blood (WHO, 2010). National and international organizations, however, have often struggled to meet these targets.

A blood shortage occurs when local blood supply fails to meet demand. This is especially prevalent in aging populations where increasing medical care is required, developing countries where chronic blood shortages are common due to limited health services and access to safe blood, and developed countries where complex blood intensive medical procedures are increasingly common (WHO, 2010). Fluctuations causing seasonal and periodic shortages are also regularly observed given the blood market operates without market prices and with short shelf-life (Slonim et al., 2014). Many regions have also experienced long-term trends like decreasing donor rates among younger populations that further contribute to low supply (Lemmens et al., 2005) (Fig 1).

**Figure 1.**
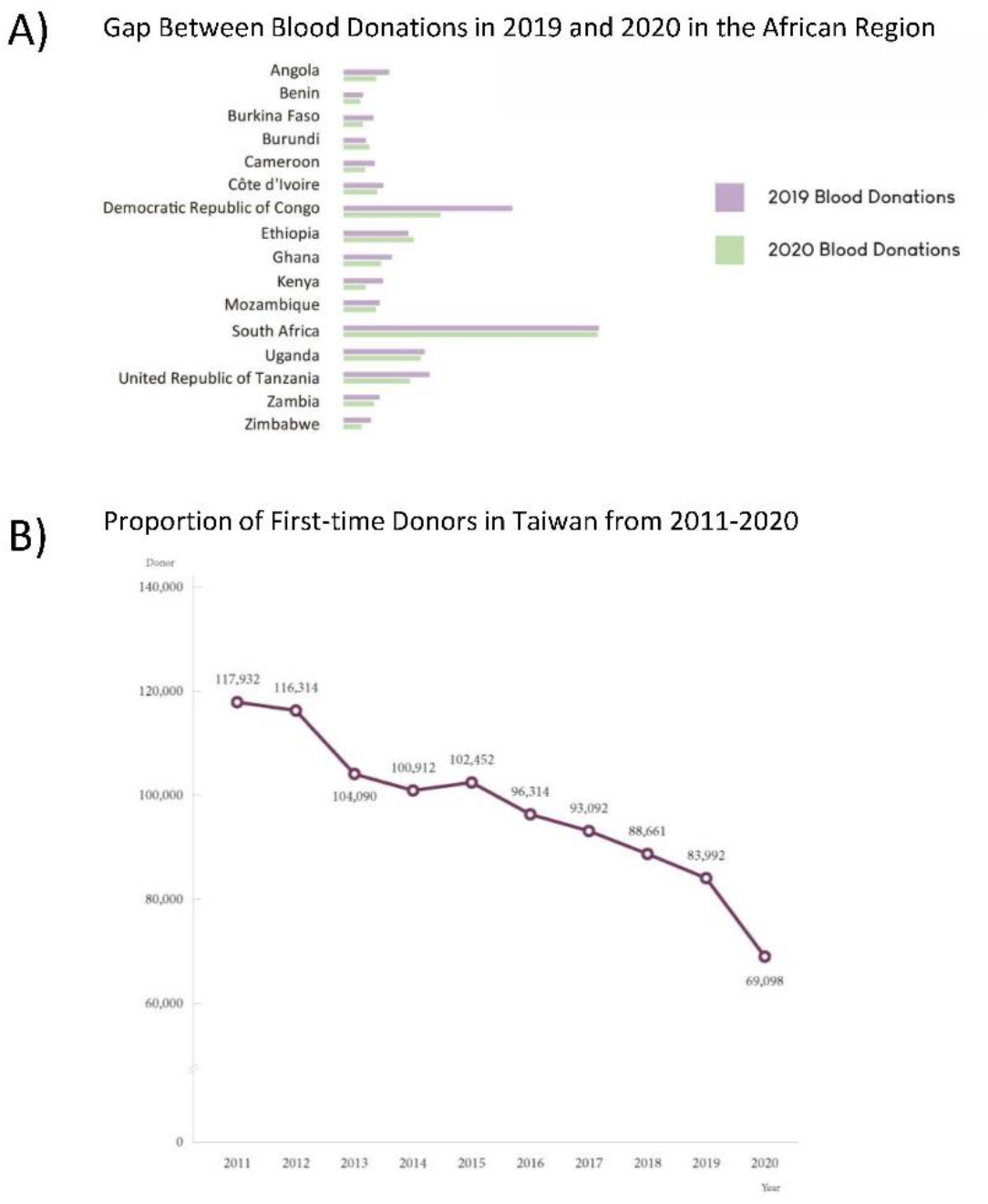
Long-term trends in blood donations show A) decreasing donor rates coinciding with the COVID-19 pandemic and B) a decreasing proportion of first-time donors that are youths (≤24 years) (Taiwan Blood Services Foundation, 2021) (Loua et al., 2021).

The COVID-19 pandemic in particular has exacerbated the blood shortage issue worldwide due to decreased blood donations and blood bank staff (Stanworth et al., 2020). Using the authors’ home residence of Taiwan during its 1st-wave surge in COVID-19 cases as an example, it has been reported that Taiwan’s blood supply has fallen to the lowest in 20 years (Taiwan News, 2021). Significant drops in donations have also been noted in the African region, and the American Red Cross reported a 10% decline in donations in the United States since the pandemic (Loua et al., 2021) (American Red Cross, 2022). The significance of blood transfusions calls for more robust measures to ensure national blood supply levels can satisfy demand.

Current attempts to alleviate the blood shortage issue include incentivization and using artificial substitutes. Some organizations have turned increasingly towards incentives such as payment, gifts, or other benefits to attract donors and increase the donor pool. Paid or commercialized donors often donate blood regularly to a blood bank for an agreed fee or sell their blood to banks and patients’ families. However, blood from these donors may not be as reliable or safe, as studies have reported that paid donors have the highest prevalence of transfusion-transmissible infections, as they may be motivated by financial factors to withhold information about their conditions. (WHO, 2010). Incentivizing and commercializing blood further raises ethical concerns as human blood donations should respect an individual’s rights to their own body and wellbeing, especially when a commercialized market may give way to exploitation (WHO, 2010).

Artificial blood substitutes are synthetically constructed to transport oxygen and carbon dioxide throughout the body. The ideal artificial blood product is safe and compatible with all blood types of the human body, is able to be sterilized to remove disease-causing viruses and microorganisms, and can be stored for over a year (Sarkar, 2008). There are currently two types of RBC substitutes being studied that differ in the way they carry oxygen: perfluorocarbon based and hemoglobin based (Moradi et al., 2016). However, problems such as solubility and stability still need to be overcome for such products to be feasible. Scientists still need to learn more about how natural RBCs function to produce a substitute with fewer side effects, increased oxygen-carrying ability, and longer lifespan in the human body (*Artificial Blood*, n.d.).

This study seeks an approach that would both preserve the altruistic nature of donations as well as the native function of red blood cells. In recent years, the notion of enzymatic blood-type conversion to eliminate transfusion compatibility has been of renewed interest (Rahfeld & Withers, 2020). The membranes of Red Blood Cells (RBCs) contain protein and sugar antigens that if recognized as foreign, elicit an adverse immune response in the human body (Dean, 2005). The presence of different types of antigens on different individuals’ RBCs is responsible for patient-donor incompatibility. While RBCs possess many different antigens, the ABO blood group system holds the greatest clinical significance in transfusions as its antigens are the most immunogenic (Mitra et al., 2014). Figure 2 depicts the basic structure of the surface antigens that distinguish A, B, AB, and O blood types. The H-antigen is a precursor of A and B antigens and is found on nearly all RBCs. The A antigen is found on type A RBCs, and the B antigen is only found on type B RBCs, while the enzymes required to create A and B antigens are codominantly expressed in AB blood types. Because type O blood lacks foreign A and B antigens, it is considered the “universal” blood type, and is often valuable for emergency transfusions and immune deficient infants, as it can be given to all patients with a Rh-positive blood type (*Blood Types Explained - A, B, AB and O* | *Red Cross Blood Services*, n.d.). Because A and B antigens on type A, B, and AB blood differ from the H-antigen only in the addition of a single sugar group, using enzymes to catalyze the cleavage of the terminal linkages on the antigens can effectively convert A and B blood to O type blood (Rahfeld and Withers, 2020).

**Figure 2.**
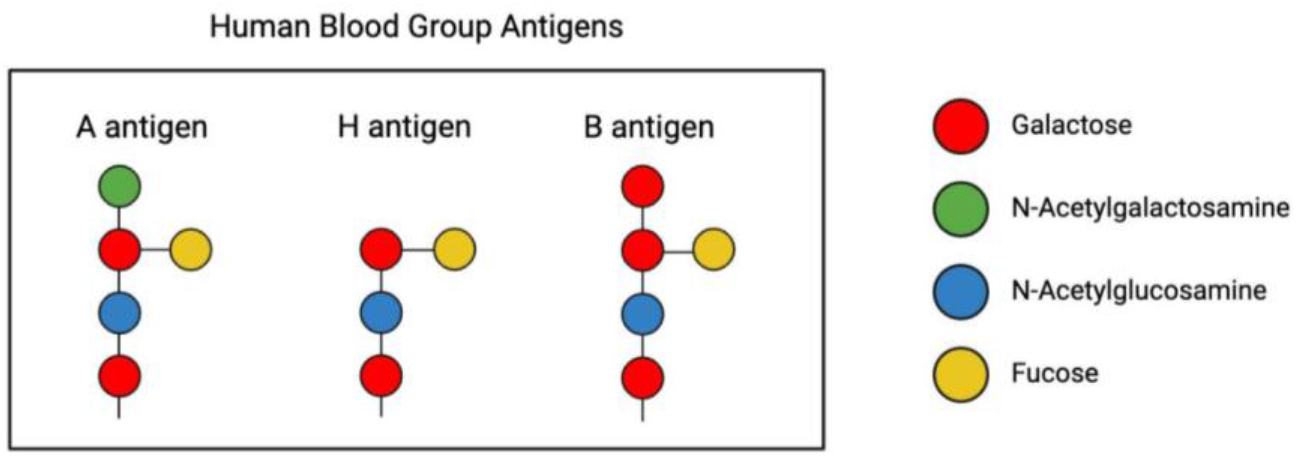
Basic Structure of A, B, and H antigens of the ABO Blood System. A and B antigens are found on Type A and B RBCs, respectively, while the H antigen is found on all RBCs, including Type O (Rahfeld and Withers, 2020).

In this study, we aim to consolidate and expand current work on enzymatic blood type conversion. Through a systematic literature search, we identified three glycoside hydrolases isolated from bacterium capable of cleaving the appropriate sites on A and B antigens to catalyze successful blood type conversion: α-N-acetylgalactosaminidase (NAGA) from *Elizabethkingia meningoseptica* that cleaves the terminal residue on A-antigens, α-galactosidase (α-gal) *Bacteroides fragilis* that cleaves the terminal residue on B-antigens, and endo-1,4-β-galactosidase (endo-β-gal) from *Streptococcus pneumoniae* that cleaves A and B antigen trisaccharides (Rahfeld & Withers, 2020) (Kwan et al., 2015) (Liu et al., 2007) (Liu et al., 2008) (Higgins et al., 2009). These enzymes were chosen based on their ability to cleave their respective blood group antigens at relatively neutral pH levels and at room temperature, properties that enable direct conversion of RBCs in blood, and eliminate the need for incubation (Table 1). By using these enzymes in combination, we hope to achieve more thorough conversion than using one enzyme alone.

**Table 1.** Properties of selected enzymes. (Rahfeld & Withers, 2020) (Kwan et al., 2015) (Liu et al., 2007) (Liu et al., 2008) (Higgins et al., 2009). Note. – The sequence for endo-β-gal used in this study was modified according to directed evolution performed by Kwan et al. The properties displayed in this table, however, reflect that of the wild type enzyme.

| | α-N-acetylgalactosaminidase<br>(NAGA) | α-galactosidase<br>(α-gal) | endo-1,4-β-galactosidase<br>(endo-β-gal) |
| --- | --- | --- | --- |
| CAZy Database<br>Enzyme Class | GH109 | GH110 | GH98 |
| Function | A-to-O conversion | B-to-O conversion | A, B conversion to O |
| Substrate<br>Specificity | Cleaves α1-3 linkage between N-acetylgalactosamine (GalNAc) residue and the β-galactose unit of the H determinant. | Cleaves α1-3 linkage between galactose residue and the β-galactose unit of the H determinant. | Cleaves Galβ-1,4-GlcNAc linkage and Galβ-1,3-GalNAc linkage that attaches A- and B-antigen trisaccharides to the rest of the antigen. |
| Source<br>Organism | *Elizabethkingia meningoseptica* | *Bacteroides fragilis* | *Streptococcus pneumoniae* |
| Functional pH | 7 | 6.8 | 8 |
| Temperature | 26°C | 25°C | 27°C |
| Molecular Mass | 50.9 kDA | 68.9 kDA | 113.8 kDA |

We aim to recombinantly produce these enzymes through high level expression in BL21(DE3) *Escherichia coli*. Then, following purification through nickel ion affinity chromatography, we evaluate the functionality of the enzymes through a combination of tests that include mass spectroscopy, which identifies compounds based on molecular mass, thin layer chromatography (TLC), which separates compounds based on polarity, and colorimetric tests, which use substrates that change color upon successful cleavage. To evaluate the ability of our enzymes to cleave actual blood antigens on the surface of RBCs, we perform agglutination tests with porcine red blood cells (pRBCs). pRBCs share a number of similar characteristics with human red blood cells and possess antigens that closely mimic the A, B and H antigens of human RBCs (Fig 3) (Smood et al., 2019) (Cooling, 2015). Because of these similarities, porcine blood is the most promising candidate for xenotransfusions, transfusions taking place across species (Long et al., 2009). The similar properties between pRBCs and human RBCs also make it an ideal substitute for human RBCs in our experiments, which allows us to avoid the safety questions that arise when working with human blood. Since the A-O antigen system exists within pRBC, the activity of NAGA can be evaluated through enzymatic reactions followed by agglutination assays with type A and O porcine blood (Mujahid & Dickert, 2015). In theory, successful cleavage of the A antigen in porcine RBC should eliminate observation of agglutination when human anti-A serum is added to the treated type A porcine blood.

**Figure 3.**
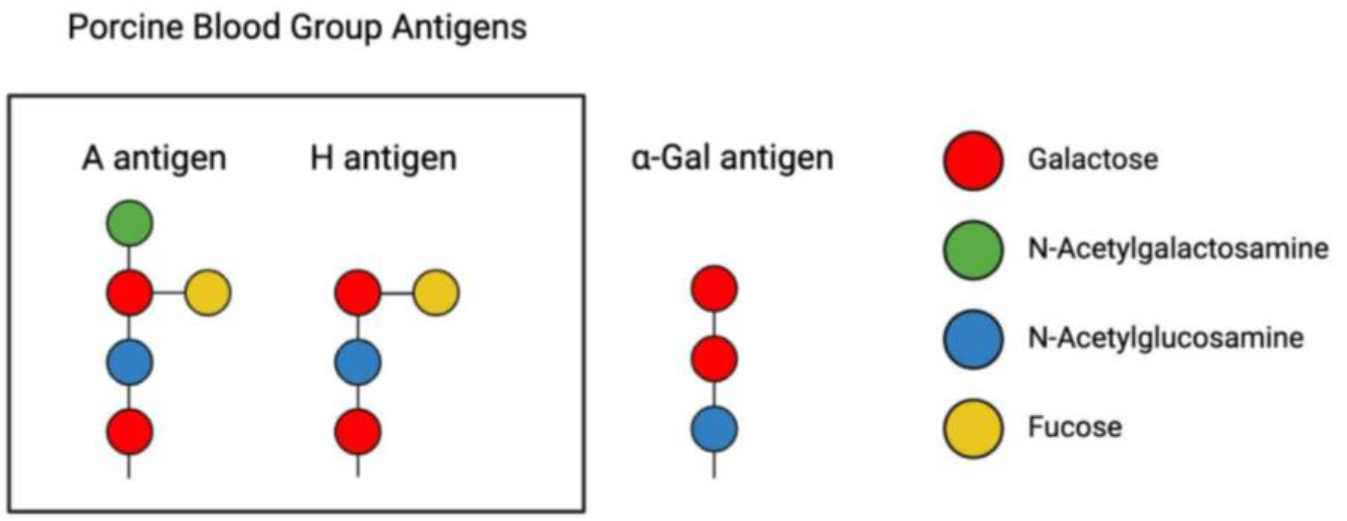
Blood group antigens found in porcine blood. Porcine RBCs have an A-O antigen system, meaning that they do not possess the B blood type antigen. Instead, porcine RBCs have a xenoantigen “α-gal,” that is present on all pRBCs, regardless of type. The structure of α-gal antigen is nearly identical to the human B antigen, but lacks a fucose group branch (Cooling, 2015).

## Materials and Methods

### Sequence Retrieval and Plasmid Design

The amino acid sequences of α-galactosidase (α-gal), α-N-acetylgalactosaminidase (NAGA), and endo-1,4-β-Galactosidase (endo-β-gal) were retrieved from the UniProt database (https://www.uniprot.org/), with accession numbers of Q5L7M8, A2AWV6, and A0A0H2UKY3, respectively. Certain amino acids of the endo-β-gal sequence were selectively modified according to a structural directed evolution performed by Kwan et al., who reported an increase in its activity towards more antigen subtypes (Kwan et al., 2015).

Following reverse translation of the sequences, gene constructs were created by attaching a T7 promoter, derived from the T7 phage and a strong ribosome binding site upstream of the enzyme sequence. A N-terminal or C-terminal 6x Histidine tag was added for protein purification purposes through a flexible Glycine-Serine linker. A double terminator downstream of the open reading frame was also added. The gene sequences were codon optimized for expression in *E. coli* using the genscript tool (https://www.genscript.com/). All composite genes were synthesized through Integrated DNA Technologies (Fig 4).

**Figure 4.**
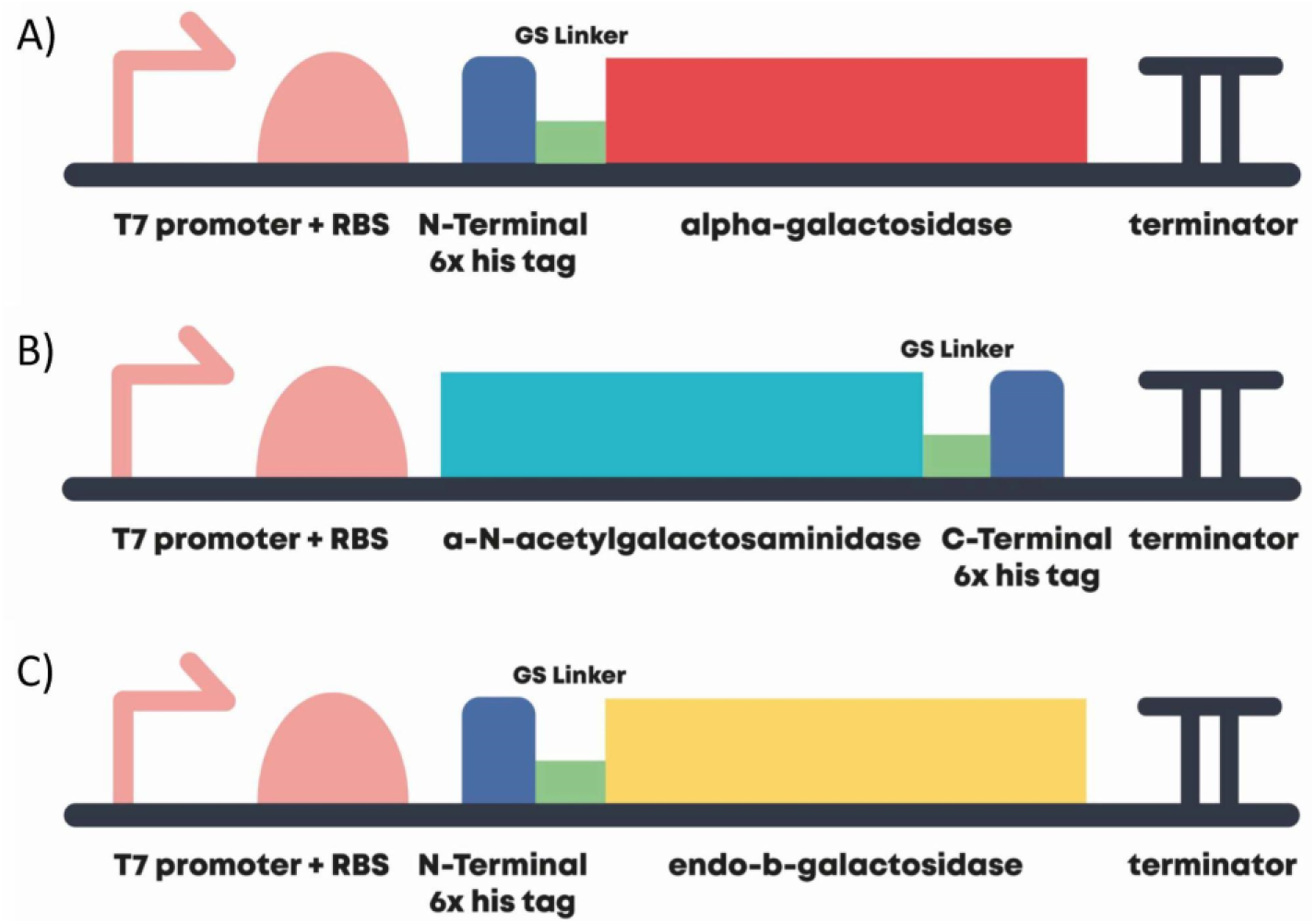
Designed construct consisting of Open Reading Frame with T7 promoter, strong RBS, N-Terminal or C-Terminal 6x Histidine-tag, and Double Terminator for A) α-gal, B) NAGA, and C) endo-β-gal.

### Protein Expression and Purification

The synthesized plasmids were transformed into BL21 (DE3) *E. coli* at 37°C overnight, diluted, and grown to OD600 0.5∼0.6 at 37°C. Cultures were induced for expression with 0.5 mM IPTG and allowed to grow overnight at room temperature. Cells were harvested by centrifugation and cell pellets were lysed through xTractor Lysis Buffer (Takara, Japan) supplemented with 20 mM imidazole. Purification of our histidine-tagged proteins was done using Ni sepharose affinity chromatography. SDS-PAGE was utilized to confirm the sizes of purified proteins. Purified enzymes were transferred to a reaction condition solution (hereinafter referred to as sodium phosphate buffer) consisting of 20mM NaH_2_PO_4_, 130mM NaCl, and 1mM DTT adjusted to pH 7 using NaOH through buffer exchange dialysis at 4°C (Hsieh et al., 2003).

### Functional Tests

For enzymes that were successfully expressed, functionality was verified using a series of experiments.

#### Colorimetric Substrates

Colorimetric substrates for NAGA and α-gal enzymes were purchased from Sigma Aldrich (Fig 5). These substrates contain a 4-nitrophenol leaving group, which turns yellow upon successful cleavage in solution. The concentration of 4-nitrophenol was quantified with absorbance at 405nm using a 96 well plate assay (Held, 2007).

**Figure 5.**
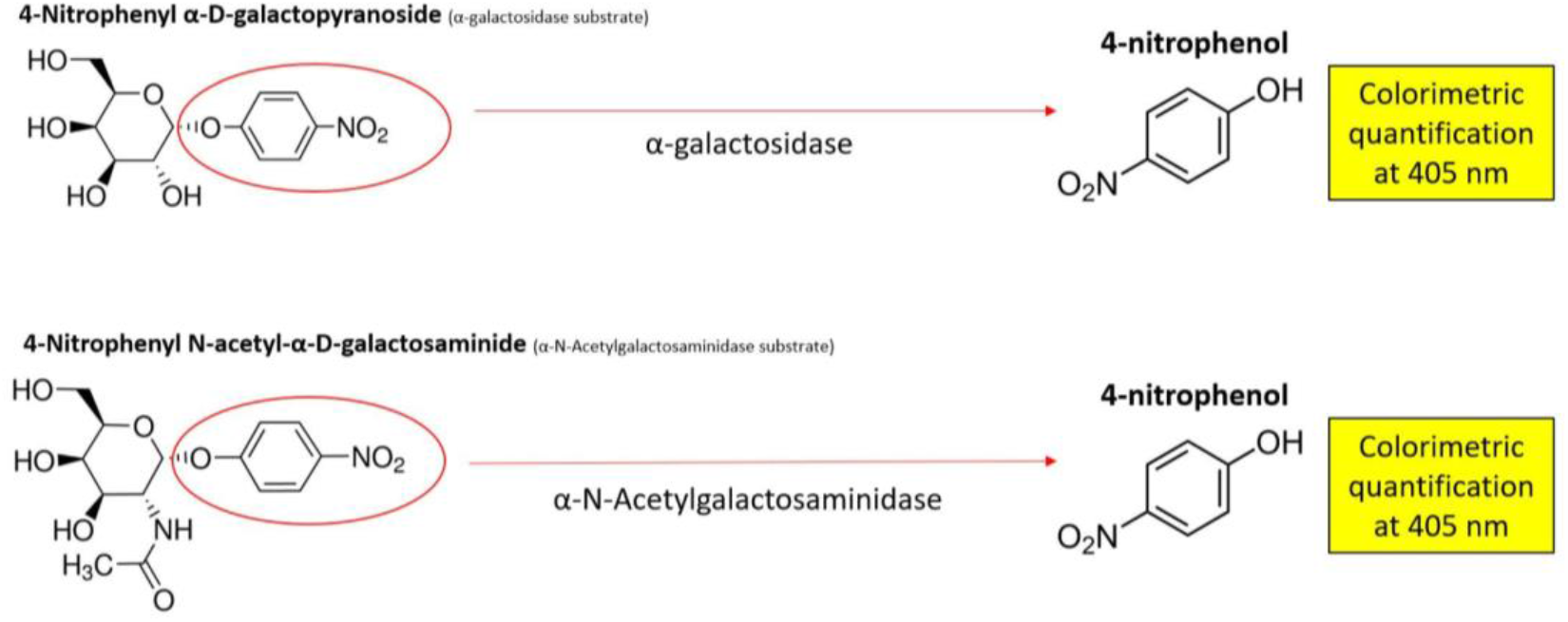
Colorimetric substrates for α-gal and NAGA. The substrates contain a 4-nitrophenol leaving group, which turns yellow upon successful cleavage in solution.

Small-scale colorimetric tests were conducted to verify the function of the enzymes. 50 μL of 10 mM substrate, 10 μL of enzyme, 30 μL of water, and 10 μL of 10x Glycobuffer 1 (50 mM CaCl2, 500 mM sodium acetate, pH 5.5), a buffer recommended by New England Biolabs to ensure optimal enzyme activity (New England Biolabs, n.d.), were added to each well (Table 2). A nonspecific substrate and nitrophenol reaction product were used as negative and positive controls, respectively. Following 2 hours of reaction at room temperature, the well contents were diluted in 1.9 mL of water and absorbance readings were taken at 405 nm with a CT-2400 Spectrophotometer (ChromTech, USA).

**Table 2.** Buffers and solutions used in substrate and control wells for colorimetric enzyme experiments.

| Substrate Well Contents |  | Control Well Contents |  |
| --- | --- | --- | --- |
| Substrate | 50 μL | Substrate (-) or reaction product (+) | 50 μL |
| Enzyme (in Sodium Phosphate Buffer) | 10 μL | Sodium Phosphate Buffer | 10 μL |
| Water | 30 μL | Water | 30 μL |
| 50 mM CaCl<sub>2</sub>, 500 mM sodium acetate | 10 μL | 50 mM CaCl<sub>2</sub>, 500 mM sodium acetate | 10 μL |

To quantify the activity of the enzymes, enzyme reactions were performed at various substrate concentrations, and the absorbance was measured at different time intervals over a constant time period with a multimode plate reader (Tecan, Switzerland). Results were averaged from at least 3 independent trials from the colorimetric assay to calculate Michaelis–Menten constants for enzyme efficiency. In order to quantitatively determine enzyme efficiency, a conversion relationship between absorbance and concentration must be determined. Therefore, a calibration curve for absorbance units versus concentration of 4-Nitrophenol product was plotted. This provided the Km values for both enzymes. With substrate concentrations equal to Km, enzyme concentrations were varied to determine the linear relationship between enzyme concentration and reaction rate. The hyperbolic relationship between substrate concentration and reaction rate for both α-galactosidase and α-N-acetylgalactosaminidase was then calculated and graphed.

#### Thin Layer Chromatography

Melibiose was used as a substrate for α-gal, followed by detection through normal-phase thin-layer chromatography (TLC), which utilizes the competition of the solute and mobile phase for binding sites to separate compounds based on polarity. α-gal can cleave the bonds in melibiose, causing the dissolution of melibiose into its monosaccharides, glucose and galactose.

To create reference standards, controls of melibiose, as well as glucose and galactose were spotted onto silica plastic-backed TLC plates (Silicycle, Canada), then ran with a mobile phase of ethyl acetate : ethanol : acetic acid : boric acid (5:2:1:1, v/v/v/v). The plate was visualized with a spraying reagent consisting of sulfuric acid : ethanol (1:1, v/v), followed by heating at 140°C for 5 minutes. Bands were observed both in ambient light and at 365 nm, and hR_f_ values were measured as the solute front divided by the solvent front multiplied by 100. These values were used as a reference when testing the functionality of α-gal. 10 μL of 10 mg/mL melibiose was incubated with 5 μL of enzyme or water (negative control) and 5 μL of 0.1 M citrate-phosphate buffer (pH 5.8) at room temperature for a total reaction time of 2 hours. The control and enzyme reaction samples were then spotted on TLC plates and developed to see if successful hydrolysis would occur, and glucose and galactose would be present in the enzyme-treated column.

#### Mass Spectroscopy

The specificity of NAGA and α-gal was further tested by using antigen trisaccharides as substrates. These trisaccharides contain three monosaccharides that have identical chemical structures to the A and B antigens found on RBCs. A reaction was carried out of the enzyme and its corresponding trisaccharide dissolved in 1X GlycoBuffer 1 (New England Biolabs, n.d.), deionized water, and purified BSA for 1 hour at 37°C. A-antigen and B-antigen trisaccharides were kindly provided by Professor Dr. Todd Lowary from the Institute of Biological Chemistry at Academic Sinica (Meloncelli & Lowary, 2010). For each enzyme, a non-specific trisaccharide served as a negative control and the trisaccharide specific to the enzyme served as the experimental unit (Table 3).

**Table 3.** Experimental setup for enzyme-antigen trisaccharide tests

| **α-N-acetylgalactosaminidase (NAGA)** |  | **α-galactosidase (α-gal)** |  |
| --- | --- | --- | --- |
| Experimental unit (+) | Negative control (-) | Experimental unit (+) | Negative control (-) |
| A-antigen trisaccharide | B-antigen trisaccharide | B-antigen trisaccharide | A-antigen trisaccharide |

After the reaction, the reaction solution was passed through C18 columns to minimize impurities from the reaction solution. The flow through solution was analyzed with a mass spectrometer to measure peaks in molar mass in order to determine the compounds present in the reaction solution. The original molar mass of the A-antigen trisaccharide is 713.35 g/mol (Meloncelli & Lowary, 2010); therefore, after cleavage, two fragments with molar masses of 510.27 g/mol and 203.08 g/mol are expected to form. The original molar mass of the B-antigen trisaccharide is 672.32 g/mol (Meloncelli & Lowary, 2010); therefore, after cleavage, two fragments with molar masses of 510.27g/mol and 162.05 g/mol are expected to form (Fig 6).

**Figure 6.**
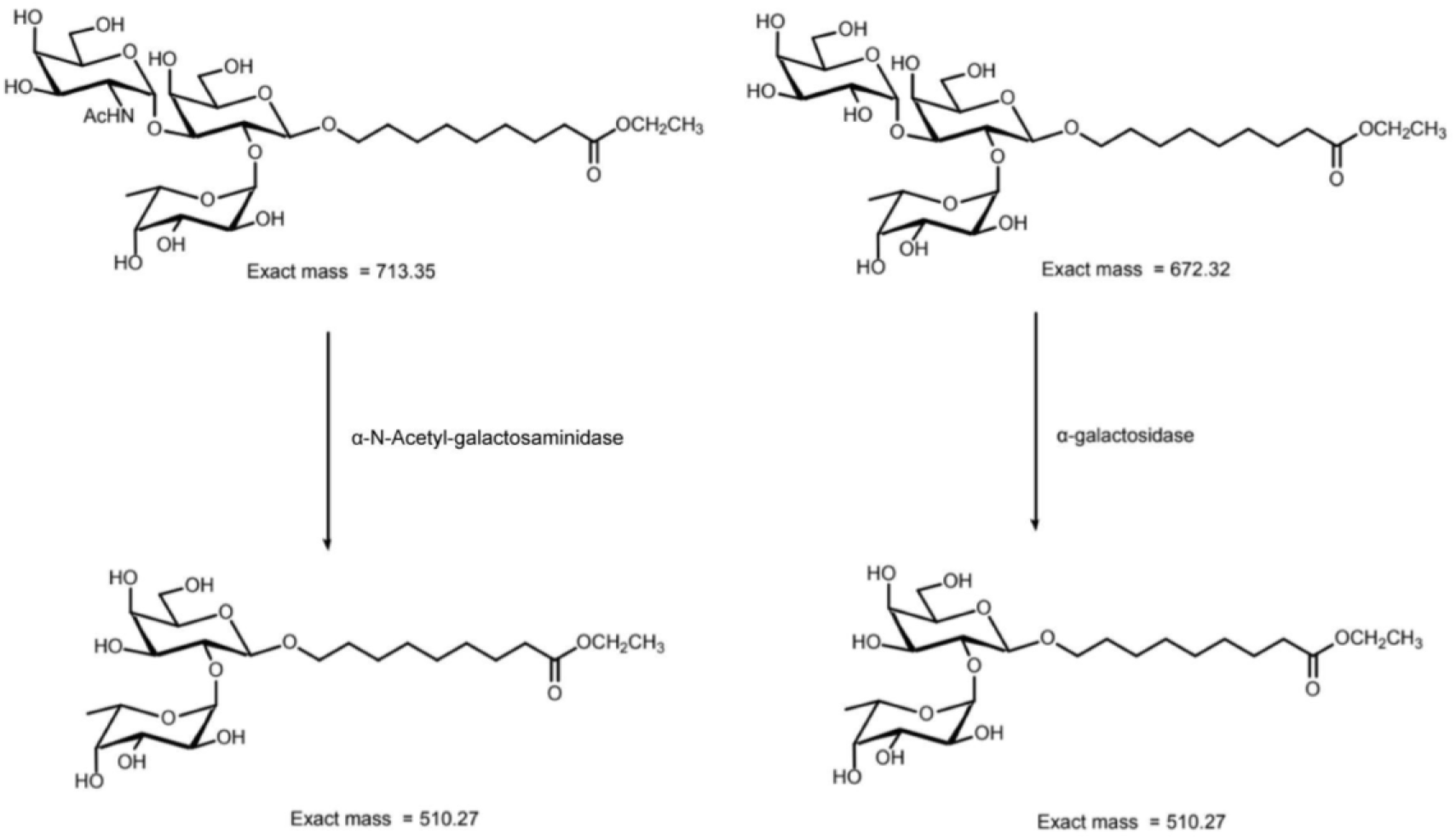
Molecular structure and mass of trisaccharides, A-antigen trisaccharide (left), B-antigen trisaccharide (right), and subsequent products following enzymatic cleavage.

### Porcine Blood Agglutination Tests

The activity of NAGA was evaluated through enzymatic reactions followed by agglutination assays with type A and O porcine blood (Mujahid & Dickert, 2015). In theory, successful cleavage of the A antigen in porcine RBC should eliminate agglutination when human anti-A serum is added to the treated type A porcine blood.

Porcine blood was sourced from a local farm in Hsinchu, a county in Taiwan, specializing in breeding pigs for scientific research purposes (https://www.atri.org.tw/). To determine the blood type of the received porcine blood, initial slide agglutination was performed with the addition of several drops of human anti-A serum to the porcine blood (Mujahid & Dickert, 2015). Previous studies have confirmed that porcine blood agglutinates well with human Anti-A antibodies, and thus should show visible clumps if the A antigen is present (Smith et al., 2006). Our results demonstrated clear clumps in some samples, and no agglutination in others (Fig 7). This difference was used to blood type our pRBCs either as A type or O type.

**Figure 7.**
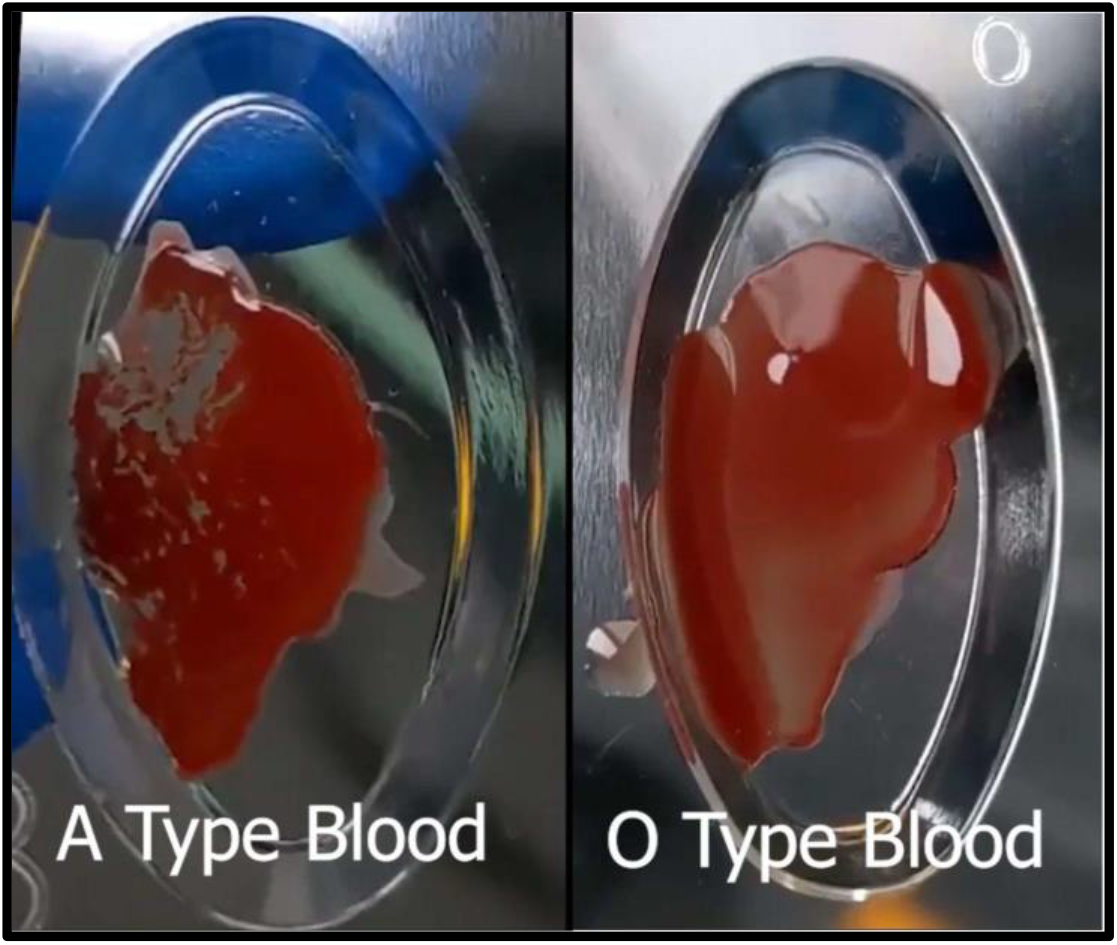
Slide agglutination of porcine blood with Anti-A serum. Clear visible clumps were observed in the left well, indicating that the sample is blood type A. There are no clumps in the middle or right well, indicating that these samples are blood type O.

To test if our NAGA enzyme works on type A pRBCs, slide agglutination was used as a qualitative test to visualize our results. Purified NAGA, previously dissolved in the sodium phosphate buffer, was dialyzed in 1X PBS to limit harm to pRBCs. Type A porcine blood seizing to agglutinate upon the addition of anti-A antibodies following treatment with the NAGA enzyme would indicate the functionality of NAGA. Non-enzyme treated type A porcine blood should continue to agglutinate as expected after the addition of anti-A serum. After enzyme treatment for 2 hours, human anti-A serum was added to the blood sample. Agglutination was visualized using 3 different qualitative methods. The blood samples and antibody mixture were spread on standard blood typing slides, to observe for macroscopic clumps, observed under a microscope slide at 1x, and under a light microscope at 20X magnification (Nikon H550S, Japan).

## Results

### SDS-PAGE confirms successful purification of recombinant α-N-acetylgalactosaminidase and α-galactosidase

Gel results show histidine-α-gal and NAGA-histidine migrating to the expected sizes of 69.7 kDa and 51.7 kDa, respectively (Figs 8-9). A clear band was observed for NAGA, while a faint band was observed for α-gal. Despite our best efforts, we were unable to express or purify histidine-endo-β-gal.

**Figure 8.**
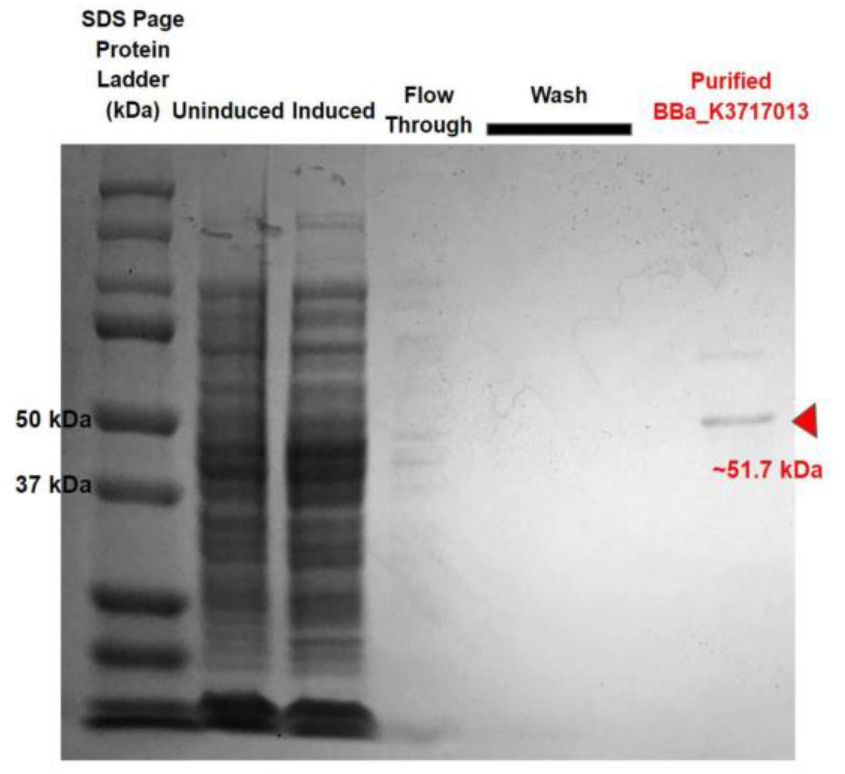
SDS-PAGE results show *E*.*coli* is able to express NAGA. The observed band is denoted by the red triangle, and compared to the ladder in the leftmost column, yields a size corresponding approximately to the expected 51.7 kDa.

**Figure 9.**
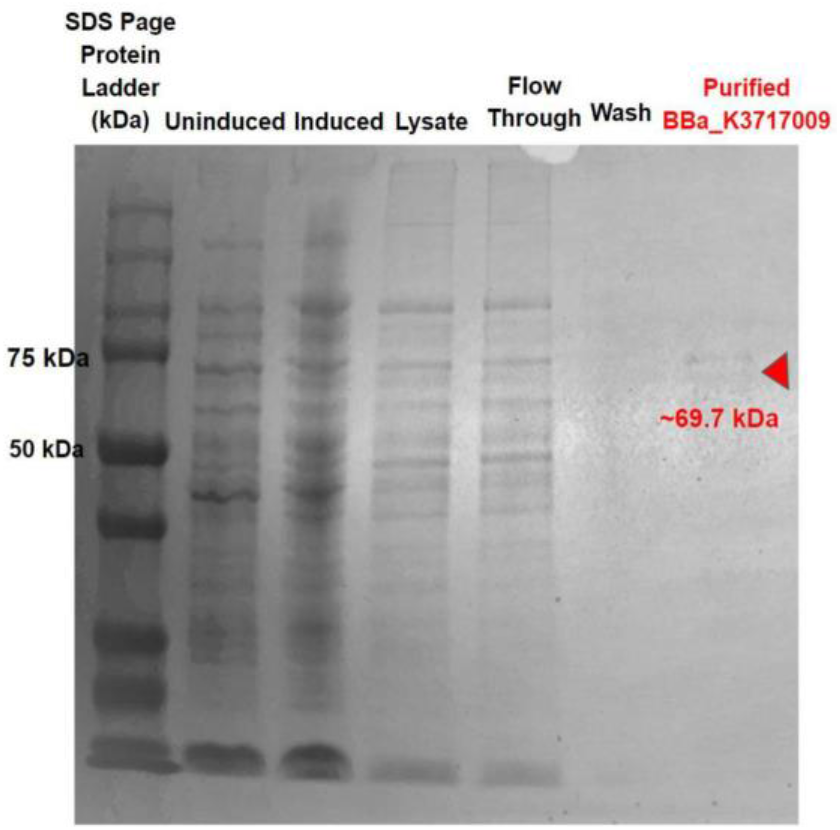
SDS-PAGE results show *E*.*coli* is able to express α-gal. The observed band is denoted by the red triangle, and compared to the ladder in the leftmost column, yields a size corresponding approximately to the expected 69.7 kDa. However, α-gal did not express as strongly as expected, producing a faint band.

### Functional tests yield corroborating results confirming enzyme activity and specificity

#### Colorimetric Substrates

Solutions with colorimetric substrates turned yellow only upon addition of the intended enzyme, demonstrating both the functionality and specificity of the α-gal and NAGA enzyme (Fig 10). When nonspecific enzymes were used in the substrate solution, the absorbance did not change, demonstrating that no cleavage had occurred. When the appropriate enzyme was added, the solution turned visibly yellow, and a high absorbance was recorded at 405 nm.

**Figure 10.**
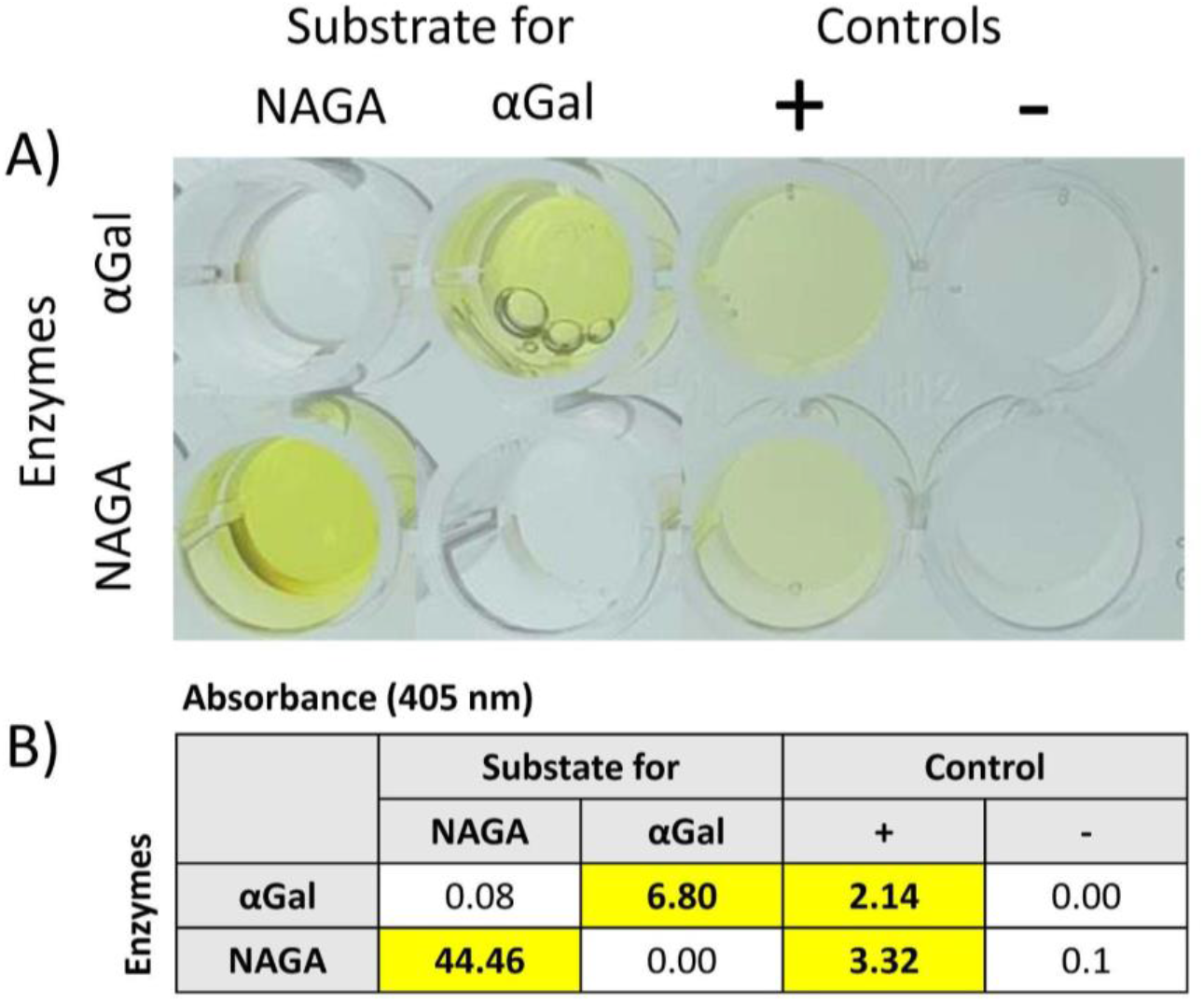
Absorbance of reaction solutions in small-scale colorimetric tests verify the functionality and specificity of the enzymes. a) 96-well plates following enzyme reaction. b) absorbance readings of diluted well contents (values accounted for dilution).

#### Thin Layer Chromatography

Based on with three independent trials with reference standards, we determined that glucose and galactose migrate significantly further up the plate compared to melibiose, with hR_f_ values of 55.9 ± 2.8 for glucose and galactose, and 33.1 ± 1.4 for melibiose (Fig 11).

**Figure 11.**
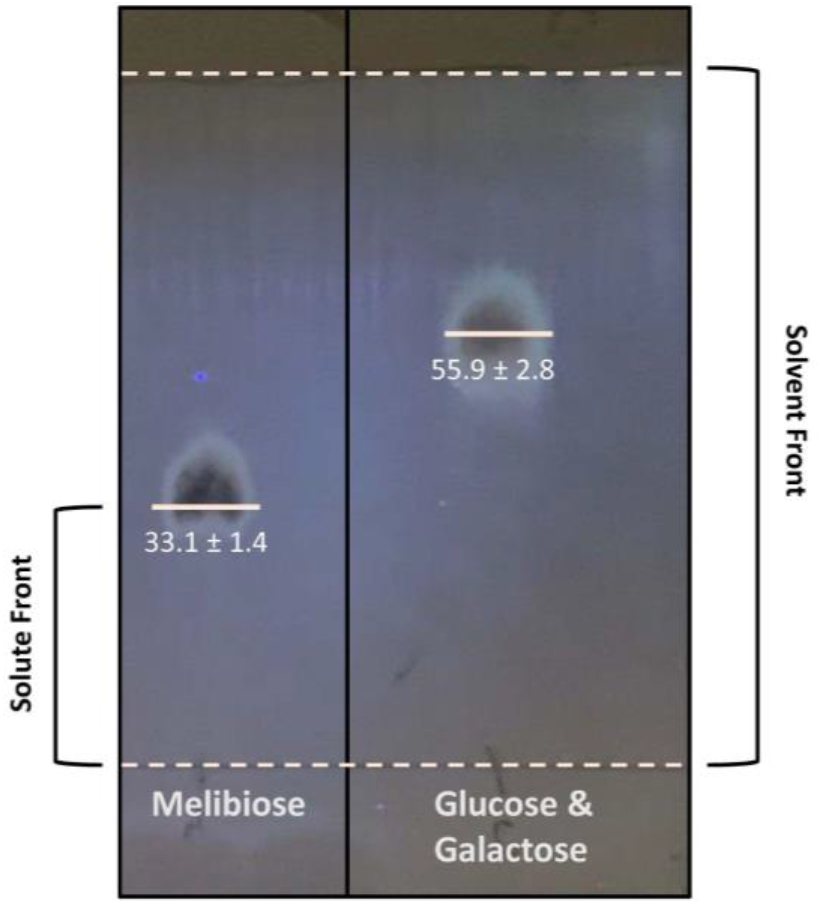
Simultaneous TLC of melibiose, glucose and galactose controls. Plate was visualized at 365 nm.

Partial cleavage was observed when the reaction product of melibiose and α-gal were run on a TLC plate (Fig 12). Compared to the control of melibiose treated with water, which only displayed one band corresponding to that of melibiose, the reaction product had two visible bands. The top band had a hR_f_ value corresponding approximately to that of glucose and galactose, indicating the enzyme had successfully catalyzed melibiose into its monosaccharide constituents.

**Figure 12.**
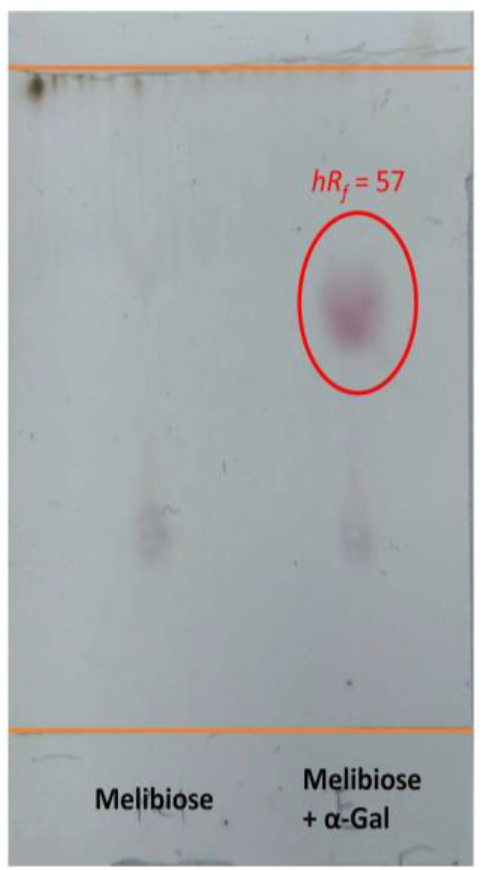
TLC plate of melibiose treated with water and α-gal under ambient light. There is clear cleavage of melibiose into glucose and galactose from the enzyme reaction, as demonstrated by the band of hR_f_ = 57 in the α-gal-treated sample, matching the predetermined value of glucose and galactose controls.

#### Mass Spectroscopy

Results obtained from the mass spectrometer further confirm the functionality of both α-gal and NAGA. For α-gal, the experimental unit shows the presence of the cleaved fragment from the reaction with the B-antigen trisaccharide, indicated by the peak at 533g/mol in the mass spectrum (Fig 13a). The negative control confirms the specificity of α-gal, as the A-antigen trisaccharide was not cleaved, indicated by the peak at 736g/mol (Fig 13b). For NAGA, the experimental unit shows the presence of the cleaved fragment from the reaction with A-antigen trisaccharide, indicated by the peak at 533g/mol in the mass spectrum (Fig 13c). The negative control confirms the specificity of NAGA, as the B-antigen trisaccharide was not cleaved, indicated by the peak at 695g/mol (Fig 13d). Throughout all readings of the mass spectrum, the molar mass of the peaks have been increased by 23 g/mol due to a sodium adduct (Kruve et al., 2013). Unrelated peaks/background noise can be attributed to compounds present in the buffer solution.

**Figure 13.**
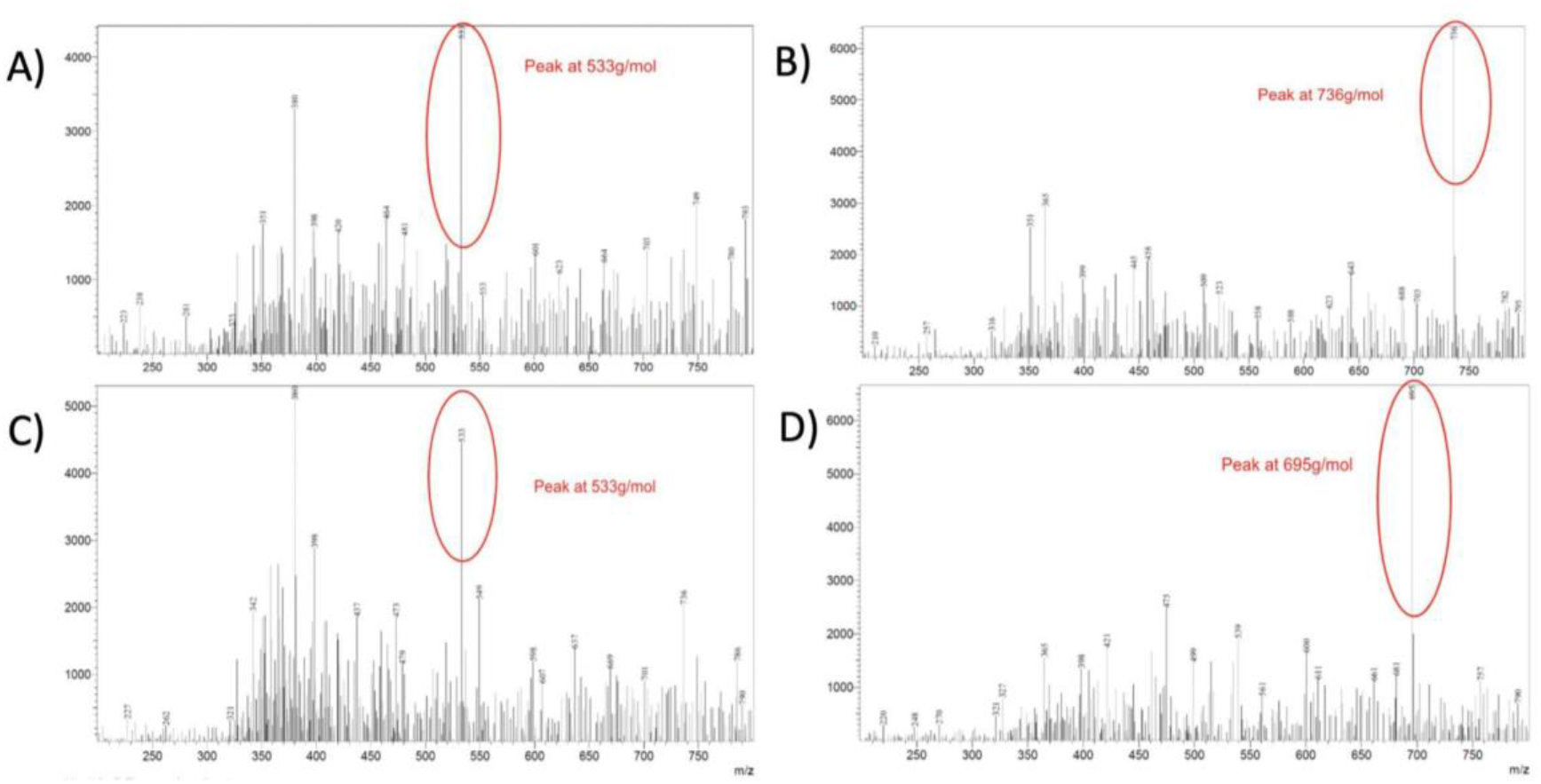
Mass spectrum of A and B antigen trisaccharide samples, with and without enzyme treatment. A) α-gal and B-antigen trisaccharide reaction solution (experimental unit, flow through). The peak at 533g/mol shows presence of the cleaved fragment from the reaction of the α-gal enzyme. B) α-gal and A-antigen trisaccharide reaction solution (negative control, flow through). C) NAGA and A-antigen trisaccharide reaction solution (experimental unit, flow through). C) The peak at 533g/mol shows the presence of the cleaved fragment from the reaction of the NAGA enzyme. D) NAGA and B-antigen trisaccharide reaction solution (negative control, flow through).

### Michaelis–Menten kinetics for enzymes were calculated based on colorimetric tests

In order to correlate the absorbance units obtained from colorimetric substrate tests with concentrations of the 4-Nitrophenol colorimetric compound, a linear calibration regression was conducted (Fig 14). The linear equation obtained was then used to convert from absorbance units to mM of 4-Nitrophenol.

**Figure 14.**
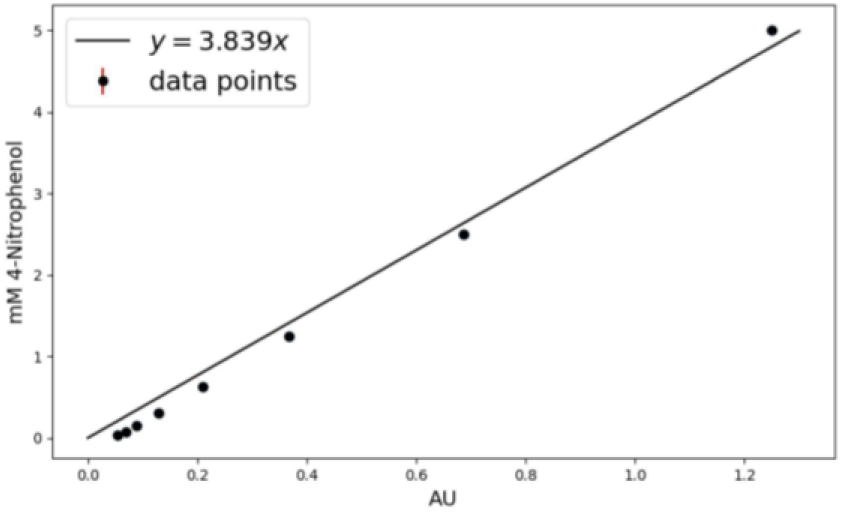
Calibration curve for the colorimetric compound (4-Nitrophenol) of the α-galactosidase and α-N-acetylgalactosaminidase substrates. Absorbance Units at 405 nm vs. mM of 4-Nitrophenol. Error bars are drawn in red but not visible due to shortness in length.

**Figure 15.**
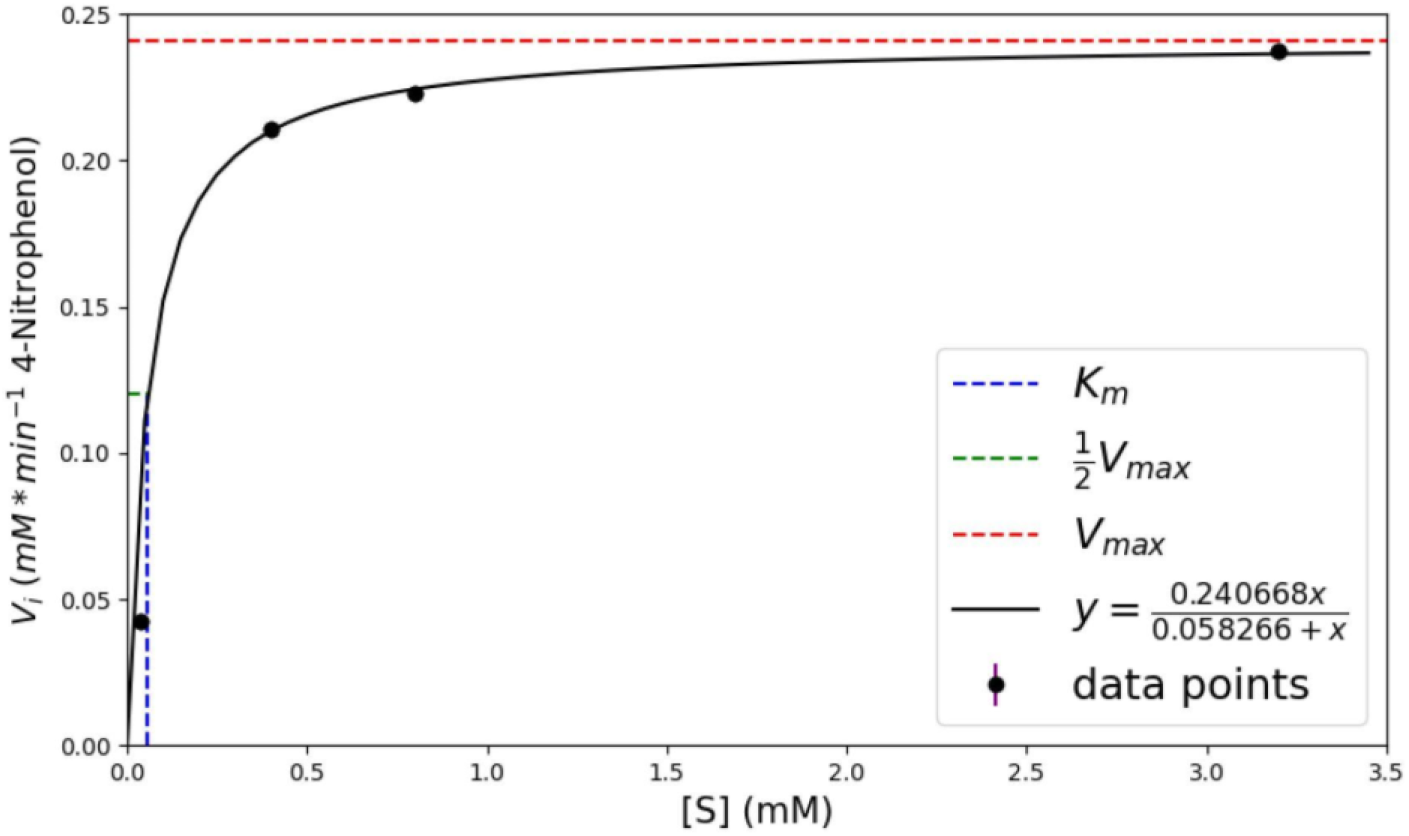
Michaelis-Menten kinetics graph for α-galactosidase. Initial Reaction Rate vs. Starting Substrate (4-Nitrophenyl α-D-galactopyranoside) Concentration Graph for α-galactosidase. Black line represents the hyperbolic equation that best fits the data points after using the Lineweaver Burk Plot. Error bars are drawn in purple but not visible due to shortness in length.

**Figure 16.**
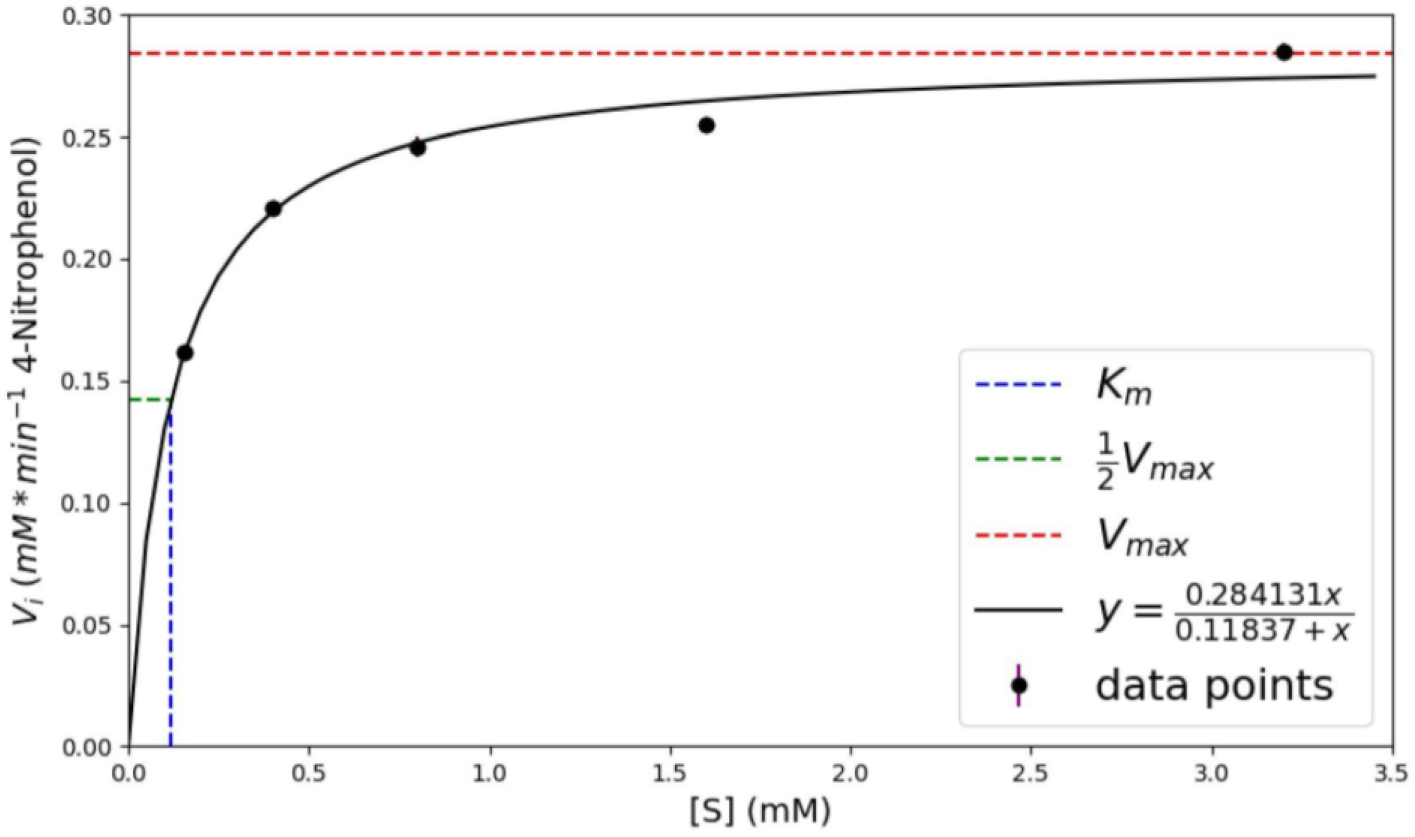
Michaelis-Menten kinetics graph for α-N-acetylgalactosaminidase. Initial Reaction Rate vs. Starting Substrate Concentration (4-Nitrophenyl N-acetyl-α-D-galactosaminide) Graph for α-N-acetylgalactosaminidase. Black line represents the hyperbolic equation that best fits the data points after using the Lineweaver Burk Plot. Error bars are drawn in purple but barely visible due to shortness in length.

To quantify the activity of the enzymes, enzyme reactions at various substrate concentrations [S] were performed and absorbance at initial intervals were measured. The slope of the initial absorbance measurements yielded the initial reaction rates Vi. At least 3 independent trials for each enzyme were used to graph and calculate Michaelis-Menten constants (Fig 17-18). As detailed in the Methods section, experiments were conducted to determine the linear relationship between enzyme concentration and maximum reaction rate (V_max_). With substrate concentration equal to K_m_ and varying enzyme concentrations, the resulting reaction rate (½ V_max_) was multiplied by 2 to obtain the enzyme concentration versus maximum reaction rate (V_max_) relationship. With substrates always in excess, these relationships (Equation 1 and 2) can help with determining the duration of reaction time and concentration of enzymes suitable for blood type conversion.

**Figure 17.**
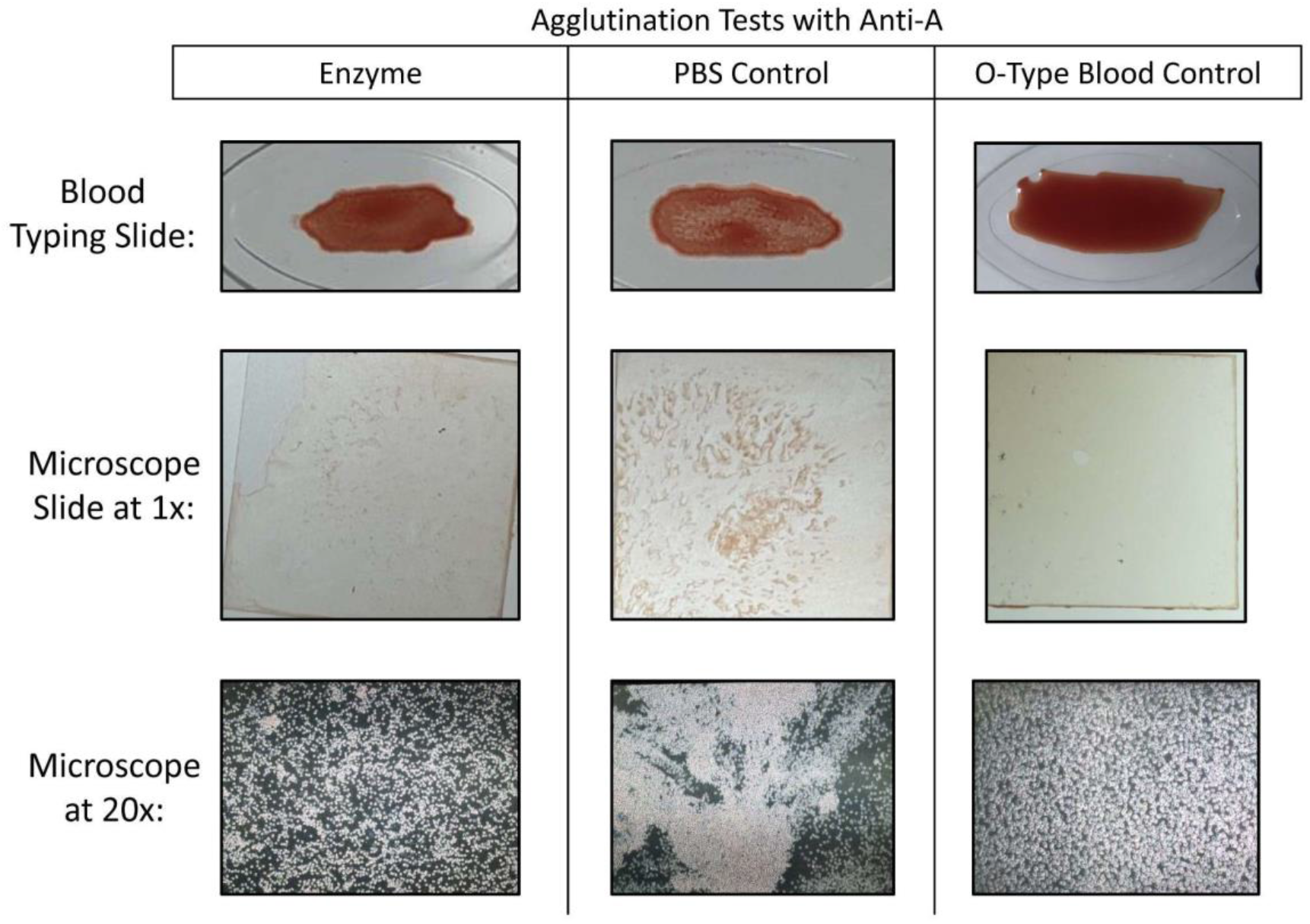
Post-enzyme treatment agglutination results visualized in blood typing wells, glass coverslips, and under light microscope at 20x. In all three methods of visualization, less agglutination was observed following enzyme treatment compared to treatment with PBS control, though there is still more agglutination compared to O-type blood. These results suggest successful A to O enzymatic conversion of pRBCs, although this conversion may not be fully complete.

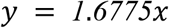

**Equation 1**. The linear function for α-galactosidase concentration (mM) and V_max_ (mM/min), with enzyme concentration being the independent variable and reaction rate being the dependent variable.

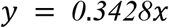

**Equation 2**. The linear function for α-N-acetylgalactosaminidase concentration (mM) and V_max_ (mM/min), with enzyme concentration being the independent variable and reaction rate being the dependent variable.

### Porcine blood slide agglutination tests demonstrate successful partial enzymatic A-to-O blood type conversion

Following enzyme treatment for 2 hours and addition of human anti-A serum to the porcine blood sample, agglutination was visualized using 3 different qualitative methods (Fig 19). First, the blood and antibody mixture was spread on standard blood typing slides, to observe for macroscopic clumps. Results showed that enzyme-treated blood displayed less clumps compared to PBS-treated blood (control), indicating less agglutination, and thus suggesting successful blood type conversion. The mixtures were then observed under a light microscope at 20X magnification. Larger and more RBC clumps in PBS control samples were observed compared to enzyme treated samples. However, enzyme treated samples still displayed more clumping than O-type blood, indicating cleavage was incomplete. The corroborative results from the macroscopic slide agglutination test and observation of RBC behavior under the light microscope provide convincing evidence of successful enzymatic cleavage by NAGA, demonstrating a proof of concept for our project.

## Discussion

Long term trends of decreasing donor rates and increasing demand as well as short term developments like the COVID-19 pandemic have exposed the susceptibility of international and national blood organizations to shortages (Lemmens et al., 2005) (Slonim et al., 2014) (Stanworth et al., 2020). The purpose of this study was to explore the method of enzymatic blood type conversion to reduce ABO antigen transfusion incompatibility, and thereby increase the supply of universally transfusable blood. Compared to alternative proposed solutions to increase blood supply, like commercialization and using artificial substitutes, such an approach has several distinct benefits. First, it avoids the ethical conflicts involved in uprooting the traditional voluntary system of blood donations. This approach also relies on converting existing RBCs rather than attempting to engineer synthetic RBCs from scratch, meaning it should be able to preserve native red blood cell function.

To evaluate this theory, we aimed to express, purify, and evaluate the functionality of 3 chosen recombinant enzymes. As mentioned, these enzymes were chosen based on their ability to cleave their respective blood group antigens at relatively neutral pH levels and at room temperature, properties that enable direct conversion of RBCs in blood, and eliminate the need for incubation (Rahfeld & Withers, 2020) (Higgins et al., 2009) (Kwan et al., 2015) (Liu et al., 2008) (Liu et al., 2007). In the first part of our project, we were able to successfully purify the NAGA and α-gal enzymes. However, despite our best efforts, we were unable to express or purify endo-β-gal. Speculations of factors that may have contributed to this include protein toxicity before induction, toxicity after induction, and the formation of inclusion bodies (Rosano, 2014). We were able to successfully demonstrate the functionality and specificity of NAGA and α-gal through a variety of tests that included TLC, colorimetric tests, as well as mass spectroscopy. We were able to verify the ability of the enzymes to cleave A-antigen and B-antigen trisaccharides in solution, demonstrating an initial proof of concept for enzymatic blood type conversion. Using porcine RBCs, we were able to successfully show an observable decrease in agglutination following NAGA treatment of A type pRBCs, suggesting partial though incomplete blood type conversion. We did not, however, notice any differences in pRBCs with and without treatment with Anti-B. We hypothesize that this is because the Anti-B serum used were ineffective in producing strong agglutination in porcine blood, and the Anti-B serum did not have the specificity to recognize the aGal antigen on porcine RBCs. To evaluate the ability of α-gal to cleave antigens on pRBCs, a different antibody agglutination mechanism with stronger and more specific antibodies may be required.

Limitations of this study include a need for more quantitative metrics that afford more sensitivity in measuring enzyme cleavage on RBCS beyond presence and absence. This will allow for the more accurate determination of the degree of enzyme cleavage when acting on actual antigens on the surface of RBCs, and eventually allow for the determination of a turnover rate of blood type conversion in a given amount of time, from which the amount of enzyme needed for conversion can be optimized. Studies have suggested using hemagglutination assays, in which samples are treated to serially diluted antibody concentrations in U and V bottom wells, and the subsequent patterns formed on the wells can be observed and assigned a standardized score corresponding to agglutination strength (Moise Jr et al., 1995). This method is also advantageous, as it sums the total scores of reaction in each well, rather than just the minimum antibody “titer” needed for agglutination (Marsh, 1972). Another more quantitative method involves using microplate technology (Mujahid & Dickert, 2015). A study has proposed a passive cost-effective microfluidic device made of filter paper, whose results can then be captured through images, and analyzed for agglutination using grayscale mean intensity through software like ImageJ, an image processing program (Samae et al., 2021). Another limitation to this study is that it does not address the immunogenic effects from other antigen systems, most notably that of the Rh blood group system. The Rh system is determined by the presence or absence of the Rhesus antigen. Because the antigen is a transmembrane protein, we believe that removal of the antigen through cleavage is implausible (Flegel, 2007). Indeed, other studies have instead opted for a “masking approach,” where polymers or single stranded DNA aptamers are attached to block anti-RhD recognition (Li et al., 2015) (Zhang et al., 2020).

In the next steps of this study, we first hope to revisit the expression of endo-β-gal. We plan to perform closer comparison of the cataloged sequences across different databases, confirm the ability of *E. coli* to express the enzyme, alter the gene sequence to reduce hydrophobicity, or utilize fusion partners to increase solubility and assist folding with chaperones if necessary. Upon successful purification of endo-β-gal, we will subject the enzyme to a similar series of functional tests such as performing TLC using lactose as a substrate (Jork et al., 1990). Next, we hope to obtain stronger and more specific antibodies, especially for human Anti-B, that will enable us to further evaluate the cleavage ability of both α-gal and NAGA on pRBCs using a hemagglutination assay as mentioned above, that will afford greater sensitivity in data collection.

## Author Contributions

All authors performed literature background research. WHT and KH designed the DNA constructs. YK, SH, WHT, JL, AC, ES, EL, and LH performed protein expression and purification. WHT, AC, and JL designed the experimental protocols. WHT, YK, SH, EL, AC, JL, LH, AL, and SK performed enzyme functional tests. SH, WHT, and AC conducted porcine blood agglutination. All authors engaged in data analysis. WHT, YK, SH, and BC drafted the manuscript. JC and JH supervised the study. All authors approved the submitted version of the manuscript.

## Acknowledgements

The authors would like to thank Dr. Todd Lowary and Louriel Macale from the Institute of Biological Chemistry at Academia Sinica for providing A and B antigen trisaccharides, as well as technical assistance with mass spectroscopy. The authors would also like to thank Mr. Sean Tsao for resource acquisition, and Dr. Nicholas Ward, for guidance on data analysis and modeling.

## Data Availability Statement

Sequences for the plasmids used in this study are available through the Registry of Standard Biological Parts. The datasets generated for this study are available on request to the corresponding author.

